# Chromatin stability generated by stochastic binding and unbinding of cross-linkers at looping sites revealed by Markov models

**DOI:** 10.1101/2020.05.22.111039

**Authors:** Andrea Papale, David Holcman

**Affiliations:** Group of Computational Biology and Applied Mathematics, Ecole Normale Supérieure, IBENS, Université PSL, 75005 Paris, France; Churchill College, University of Cambridge, CB30DS, UK

## Abstract

Chromatin loops inside the nucleus can be stable for a very long time, which remains poorly understood. Such a time is crucial for chromatin organization maintenance and stability. We explore here several physical scenarios, where loop maintenance is due to diffusing cross-linkers such as cohesin and CTCF that can bind and unbind at the base of chromatin loops. Using a Markov chain approach to coarse-grain the binding and unbinding, we consider that a stable loop disappears when the last cross-linker (CTCF or cohesin molecule) is unbound. We derive expressions for this last passage times that we use to quantify the loop stability for various value parameters, such as the chemical rate constant or the number of cross-linkers. The present analysis suggests that this binding and unbinding mechanism is sufficient to guarantee that there are always cross-linkers in place because they generate a positive feed-back mechanism that stabilizes the loop over long-time. To conclude, we propose that tens to hundreds cross-linkers per loop are sufficient to guarantee the loop stability in the genome over a cell cycle.

## INTRODUCTION

Chromatin organization in the nucleus of eukaryotic cells remains largely unclear, but several clues can be revealed by the high-resolution Hi-C experiments [1] that provide the distribution and contact probabilities of genomic pairs. In particular, this analysis has revealed that chromatin organization is characterized by three dimensional structures enriched in loops, called Topologically Associating Domains (TADs). These TADs appear as bloc-submatrices in the overall contact matrix. From a genomic viewpoint, TADs constitute regulatory neighborhoods that participate to isolate genes and their associated regulatory elements from the rest of the genome. Interestingly, one of the role of TAD structures is to prevent the formation of inappropriate gene-enhancer contacts between neighboring domains, as TAD disruption can induce carcinogenesis [2, 3]. These TADs are formed by the chromatin itself with highly enriched intra-domain interactions [4] and are described in polymer physics as an ensemble of short and long-range loops [5]. These loops are controlled by nanodomains enriched in specific binders or cross-linkers [6, 7] such as cohesin and CTCF proteins that bind to the majority of borders that demarcate TADs [8]. CTCF binding sites are among the most frequent mutation/deletion hotspots in multiple cancer types and these mutations/deletions at TAD borders can activate oncogenes [3].

Our aim in this manuscript is to explore how already formed loops can be maintained for long time. Indeed, although recent studies focused on loop formation [9, 10], very little is known about the possible mechanism that can maintain loops for long time. To study the loop stability, we develop a Markov model of binding and unbinding for cross-linkers at the base of a formed loop. Indeed, loop formation experiments [9, 10] suggests that loop-extruding complexes contain more than one molecules. In our the present model, we suppose that the initial state includes the presence of a cohesin ring, embracing two strands of chromatin, which has already reached two CTCF binding sites (with opposite orientations), both of them occupied by at least one CTCF molecule. We assume that when the last CTCF molecule leaves the base of a loop, the loop disappears. We will derive a last passage time formula, using a Markov-chain modeling approach with specific absorbing boundary conditions [11, 12] (reviewed in [13–15]). We will then use our analytical formulas to explore chromatin loop stability. The present analysis reveals that these loops can be stabilized for long-time, in agreement with the presence of stable chromatin regions required to express specific genes and repress others.

The manuscript is organized as follow: in the first part, we introduce the physical model of binding and unbinding cross-linkers to the loop nanoregion. In the second part, we analyze three possible scenarios of cross-linker stabilization. In the third part, we report some finding and in particular, that the mechanism of binding, occurring preferentially at the center of a looping region by a dynamical equilibrium, can generate a long-term stability with few cross-linkers, which could explain the inherent long-term memory of the genome organization before cell division.

### A MARKOV MODEL OF CROSS-LINKER BINDING TO A NANODOMAIN LOOPING REGION

#### Case 1: Uniform binding and unbinding rates of cross-linkers

We start by describing the physical model we use here to study the interactions of diffusing cross-linkers with a nanoregion at the base of a formed loop. We use a Markov chain model to coarse-grain the motion of *N* diffusing cross-linkers inside the nucleus, modeled as a sphere. Cross-linkers diffuse until they enter a nanodomain region Ω (orange in Fig.1), which represents a binding region at the base of a loop. As long as there are enough cross-linkers in this region, we consider the loop stable. Outside Ω, the cross-linker molecules diffuse freely. We coarse-grain the diffusion flux into Ω as an inward rate *λ*. We do not limit the number of cross-linkers in such region. Once a cross-linker enters Ω, it can stay there for a random time, which is Poissonian with rate *μ*.

**FIG. 1.**
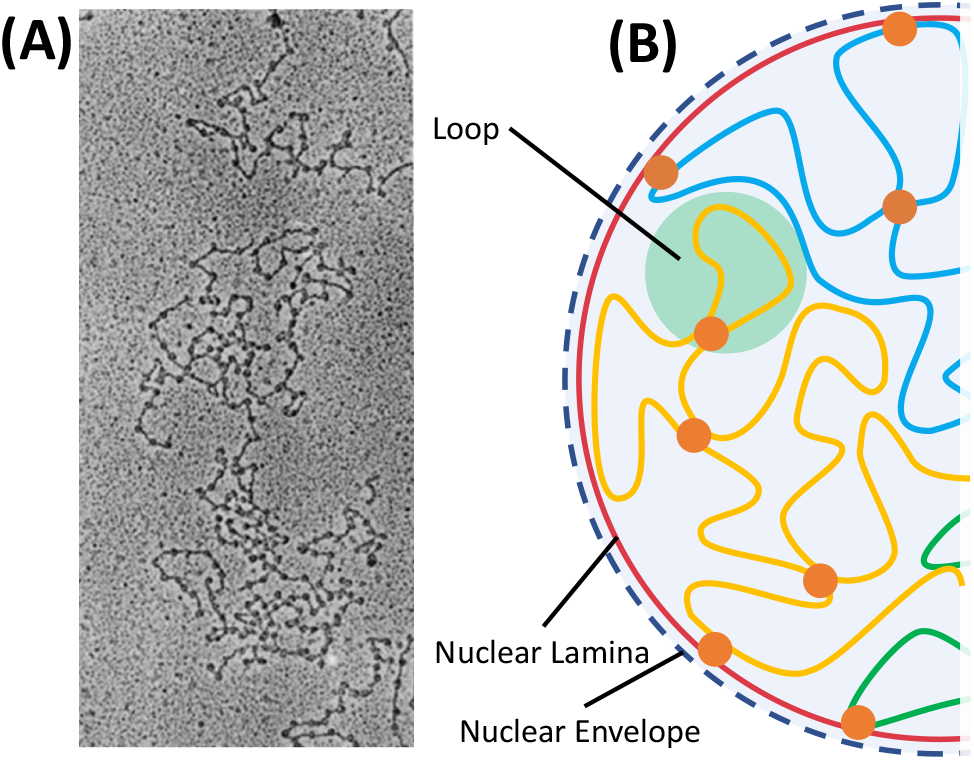
Illustration of chromatin Loops. (A) Electron-microscopy images of DNA showing many loops. (B) schematic representation of chromatin inside the nucleus ith its envelop (red and dashed blue lines), where cohesin is marked (orange dots) and often present at the base of a loop. Image reproduced by [16].

We shall evaluate three scenarios of escape when the cross-linkers: 1- can escape independently (Fig.2A), 2- only at the two extremities (when the cross-linkers are organized linearly) can escape (Fig.2B), 3- escape with a rate which is rescaled by a factor *p* ∈ [0, 1] except for the two extremities (Fig.2C).

**FIG. 2.**
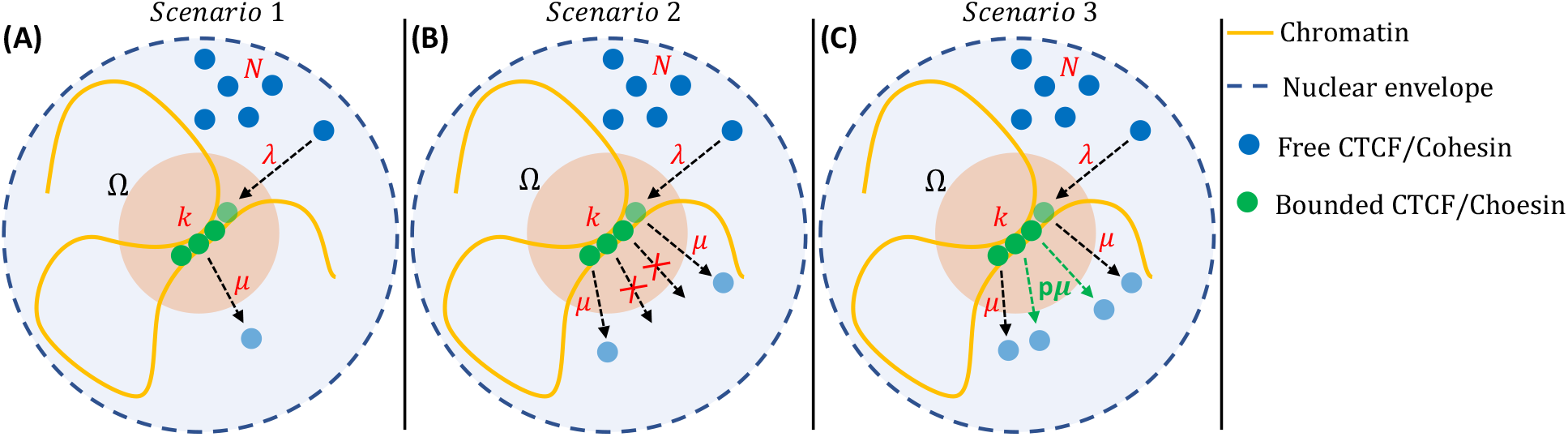
Three scenario for cross-linker bindings in a nanoregion to maintain chromatin loops. (A) The escape rates *λ* (binding) and *μ* (binding) do not dependent on the number of cross-linkers; (B) the cross-linkers are organized linearly and only the two extremes can escape; (C) cross-linkers are still organized linearly but the escape rate *λ* is rescaled by a factor *p* (0 ≤ *p* ≤ 1) except for the two extremes case.

Our goal is to estimate the number of cross-linkers inside Ω. As long as Ω contains at least one cross-linker, the loop persists. When the last one escapes, the loop is lost and the region Ω disappears as well as the memory of the local chromatin organization. Thus, to measure the loop stability, we compute the mean first passage time for the last cross-linkers to escape. To compute such time, we follow the number *n* (*t*) of particles trapped within Ω at time *t* (only one particle at a time can enter) [12]. The probability density function (pdf) that there are *n* (*t*) = *k* cross-linkers at time t inside Ω is for 0 ≤ *k* ≤ *N*,

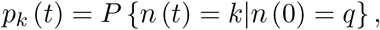

when there are initially *n* (0) = *q* cross-linkers inside Ω. The transitions to the state where there are exactly *k* cross-linkers, is computed by considering the changes during time *t* and *t* + Δ*t*:

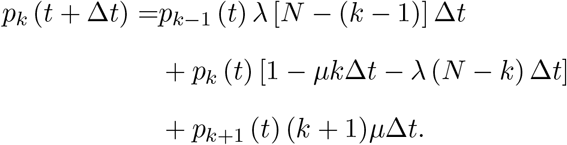

For *k* = 0 and *k* = *N*, we get

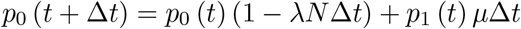

and

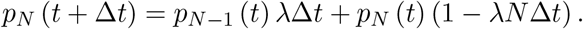

Thus, we obtain a Markov system of *N* + 1 equations

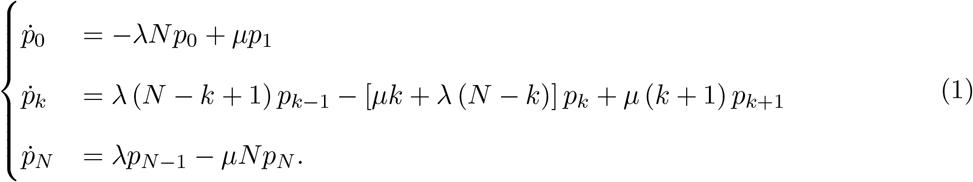

To compute the first time that all cross-linkers escape the region Ω, the state *k* = 0 should be absorbing so that the Markov chain is now modified to

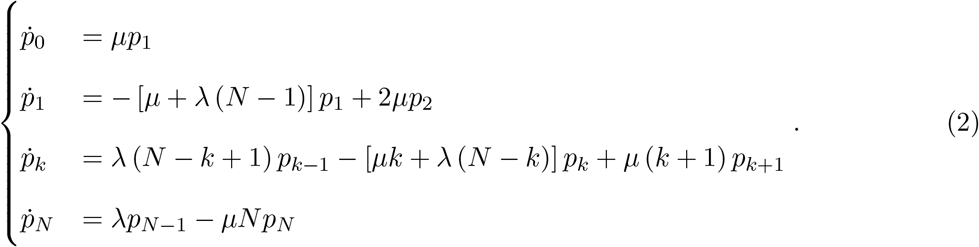

The *mean time to lose a loop* 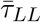 is the first time when all cross-linkers have left the nanoregion. We refine the definition by considering th time 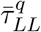 when there are initially *q* bound cross-linkers:

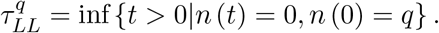

Note that the probability of the time random time 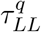 is link to the Markov chain by the relation

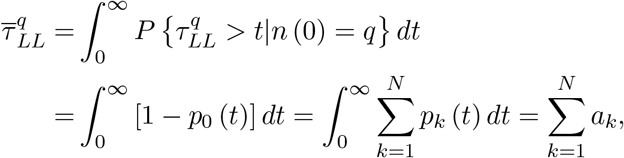

where

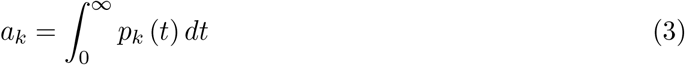

and the normalization condition is 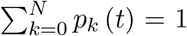. We thus need to compute the *a*_*k*_ coefficients (see Appendix). We obtain using an induction procedure that

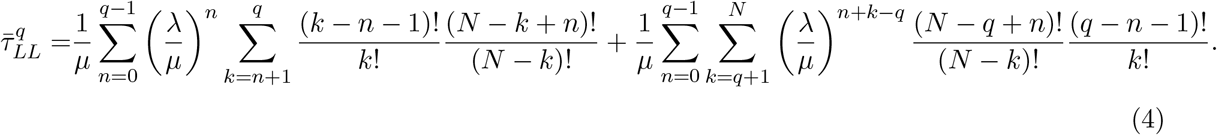

In the limit case where *λ/μ* ≪ 1, this expression can be approximated at first order using 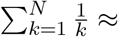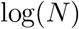 and 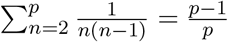 by

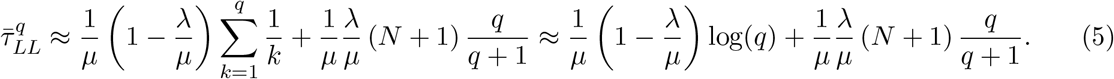

In the opposite limit *λ/μ* ≫ 1, we obtain using the leading order term,

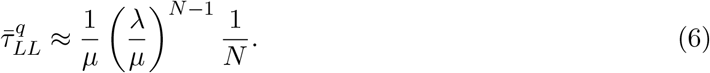

We plotted the time 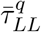 with respect to various parameters in Figs 3–6 and we will discuss the results below in comparison with the two other models.

**FIG. 3.**
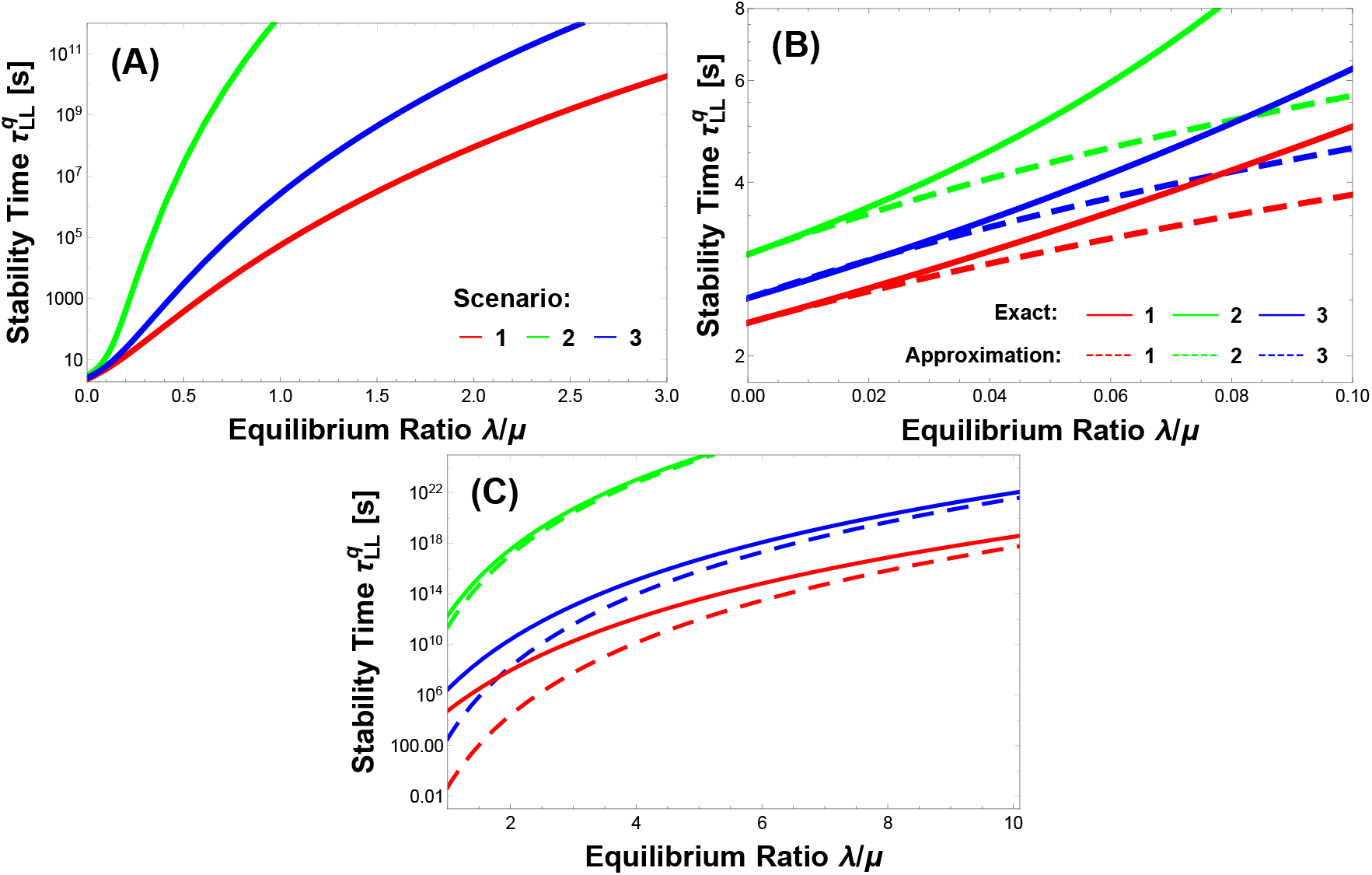
Stability time 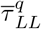 vs equilibrium ratio *λ/μ* for the three scenario. (A) Analytic expressions in the three scenario considered. (B) Limit *λ/μ* ≪ 1, comparison of the exact relations (continuous lines) with the approximated equations 5, 9 and 12 (dashed). (C) Limit *λ/μ* ≫ 1, comparison (continuous lines) with the approximated equations 6, 10 and 13 (dashed). Total number of *N* = 20 when there are *q* = 5 molecules initially. The forward rate is set to *μ* = 1 *s*^−1^. For the third scenario, 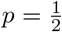.

#### Case 2: Unbinding rate depends on 2 cross-linker and the binding rate is uniform

We consider here the case where the unbinding can occur through the two external cross-linkers organized in line (Fig. 2B). The two extreme cross-linkers can escape with a rate *λ*, while the others have to wait until their turn. The probability to find *k* particles is now

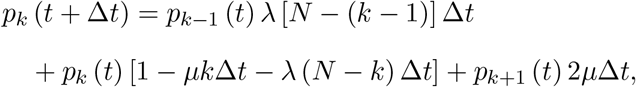

and the new equations are (state 0 is absorbing):

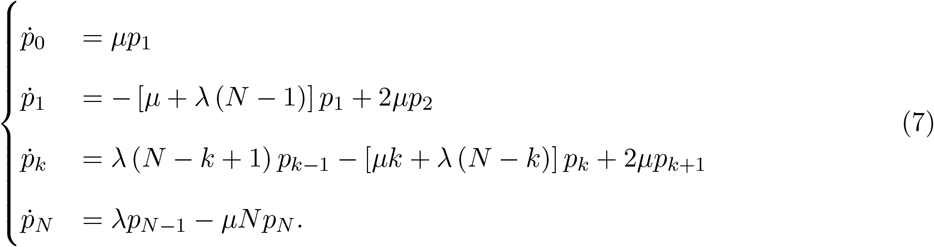

Following the method of induction as described in Appendix (see Appendix), we obtain the general expression:

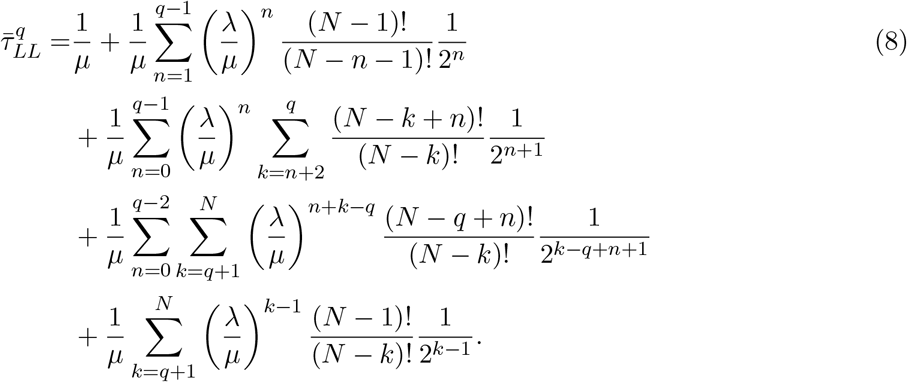

In the limit *λ/μ* ≪ 1,we approximated this sum using the first order in *μ/λ*. With 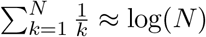 and 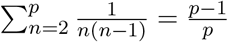 we get:

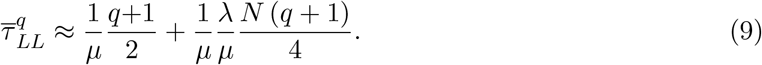

In the opposite limit *λ/μ* ≫ 1, this sum can be approximated at the last order of *λ/μ*, leading to

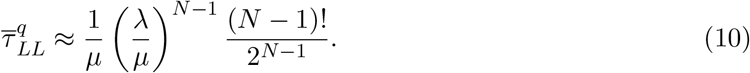

#### Case 3: Non-uniform unbinding rate *λ* and uniform rate *μ*

In this case, we consider here that the two external cross-linkers can escape with a rate *λ* while the others located inside (Fig. 5) can detache and escape with a modified rate *pλ* where *p* ∈ [0, 1] is a free parameter. The probability to find *k* particles in the domain Ω, for *k* > 2 is now

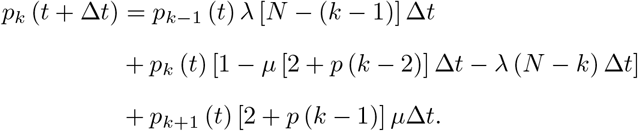

The Markov chain where the 0-state remains absorbing is

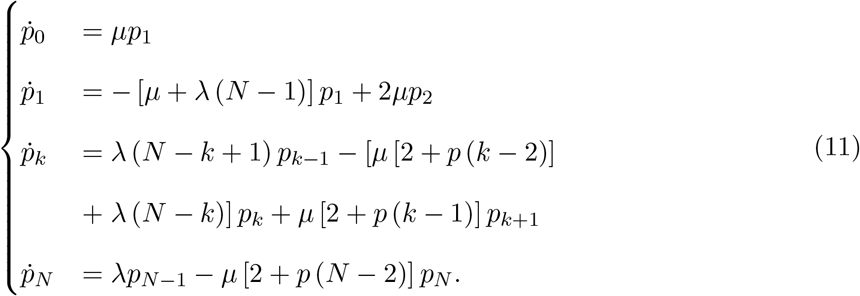

Finally, we obtain the general expression using the method of induction as described in Appendix (see Appendix):

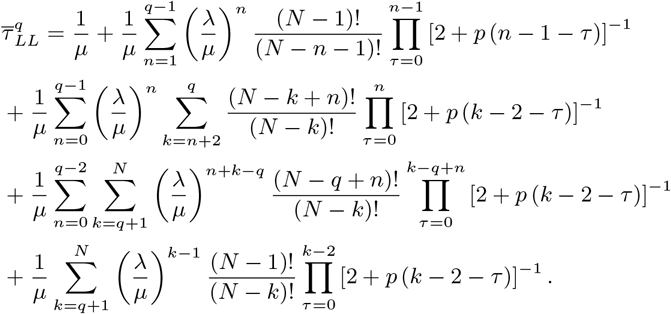

In the limit *λ/μ* ≪ 1, this sum can be approximated at the first order of *μ/λ*. Using 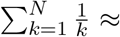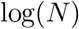 and 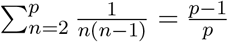 we get:

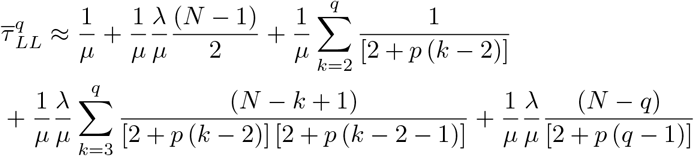

By expanding 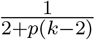 as 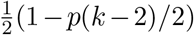 and 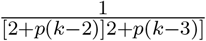 as 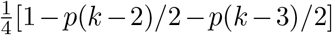, we get

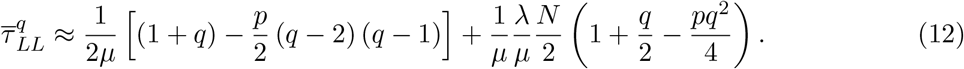

In the limit *λ/μ* ≫ 1, 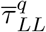 can be approximated at the last order of *λ/μ*:

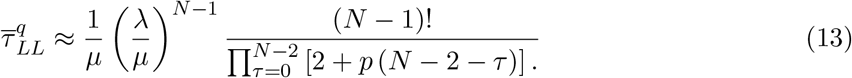

#### Sensitivity analysis of the residence time 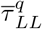 with respect to the main parameters

We now compare the results for the residence time 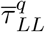 for the three scenario. As we shall see, the time of loop maintenance by the dynamics binding is quite robust. We first plotted the stability time 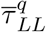 vs equilibrium ratio *λ/μ* for the three scenario in Fig. 3A and found that for a small variation from 1 to 3 the time increases from order of magnitude, demonstrating the robustness of the process. We also compare the approximation to the full solution in Fig. 3B-C.

To further explore the stability of the time 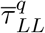, we vary the total number of cross-linkers and found again a robust time when N varied from 5 to 30 in all scenarios (Fig. 4). Finally for scenario 3, we varied the releasing parameter *p* (*p* = 1 recovers the first scenario while *p* = 0 corresponds to second). Finally, we vary the initial number *q* of bound cross-linkers (Fig. 6): except for a small *λ/μ* ratio, the time increases extremely quickly independent of the initial values q.

**FIG. 4.**
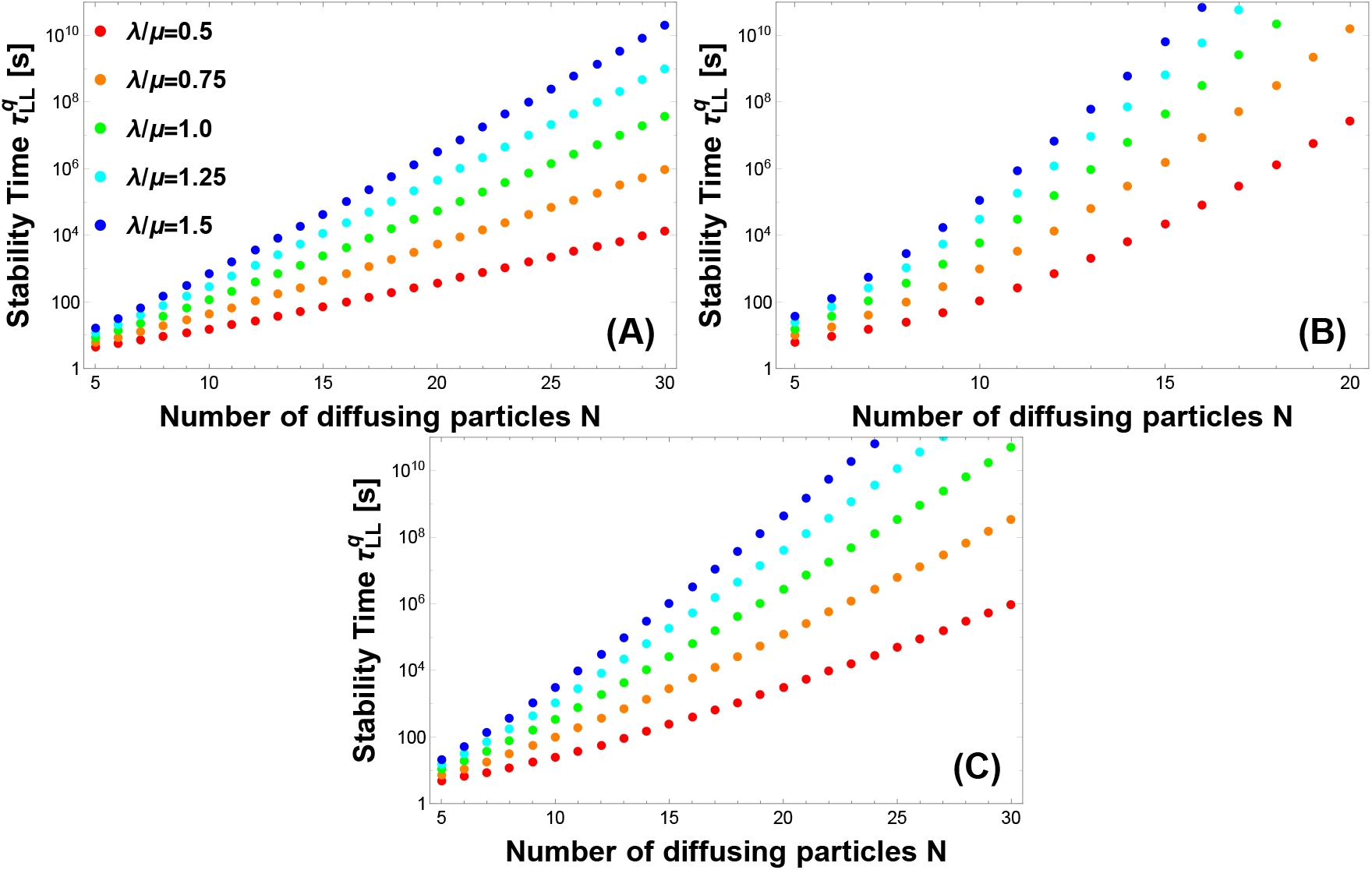
Stability time 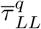 vs the total number of cross-linkers *N* for the three scenarios considered. (A) scenario 1; (B) scenario 2; (C) scenario 3. There are *q* = 5 molecules initially. The forward rate is set to *μ* = 1 *s*^−1^. For the third scenario, 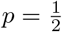.

**FIG. 5.**
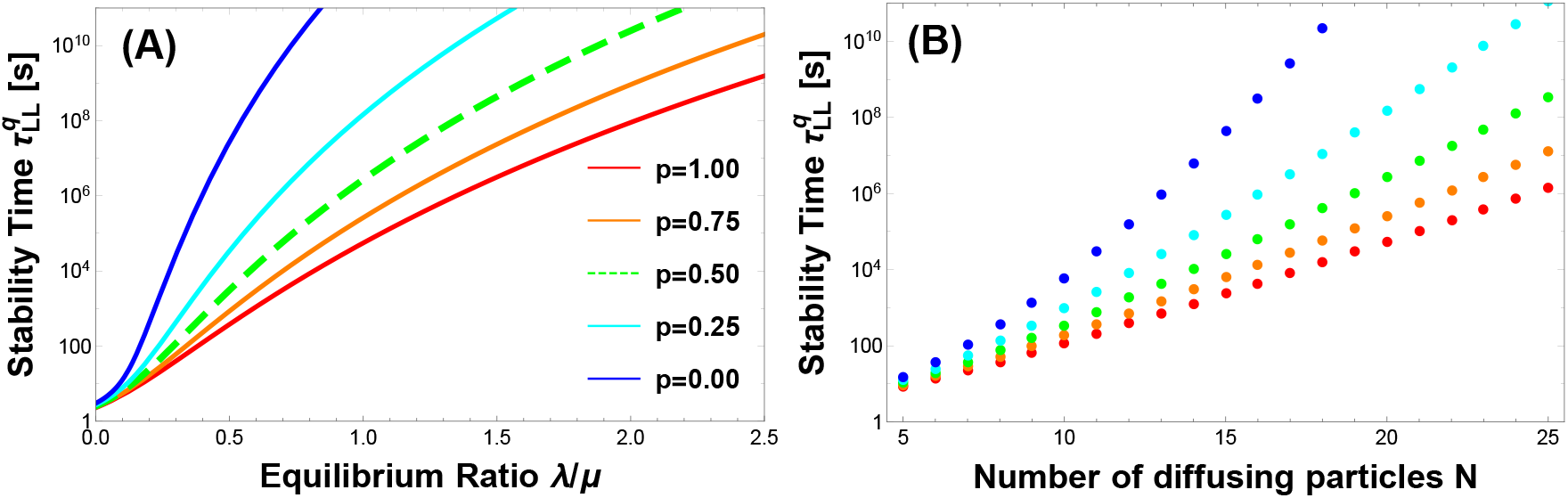
Effect of the releasing parameter *p* for the third scenario. Stability time 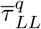 vs *N* (A) and equilibrium ratio *λ/μ* (panel B, where *N* = 20) for various values of the parameter *p*. There are *q* = 5 molecules initially and the forward rate is set to *μ* = 1 *s*^−1^.

**FIG. 6.**
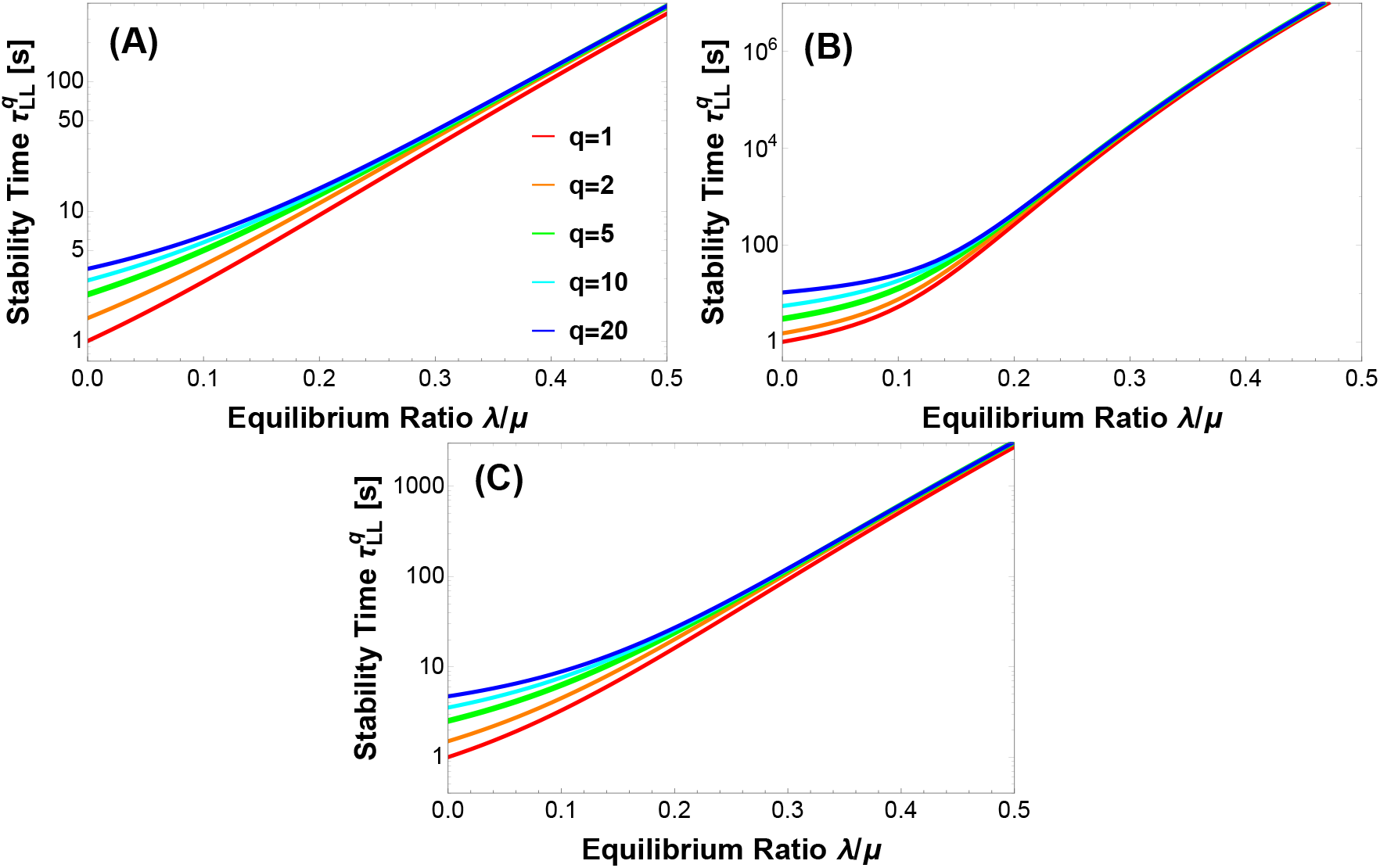
Effect of changing the initial number of bound cross-linkers on the stability time 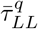 for the three scenario. (A) scenario 1; (B) scenario 2; (C) scenario 3. Total number of *N* = 20 where there are *q* = 5 molecules initially. The forward rate is set to *μ* = 1 *s*^−1^ For the third scenario, 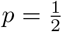.

## DISCUSSION AND CONCLUDING REMARKS

We have presented three scenarios based on a rate model to study the stability of already formed chromatin loops. The underlying model assumes that cross-linkers bind to a nanodomain region to stabilize a loop. Although these scenarios differ by the unbinding rates (uniform, unbinding from the extreme when organized linearly and partial unbinding from all cross-linkers), they all converge to show that few bound molecules in a neighborhood of the looping closing region are sufficient to guarantee that the number of cross-linkers will maintain the loop for a long-time. Interestingly, the number is around 10 (resp 15) cross-linkers per loop, we found that a loop can be maintained for days (resp. months).

When transcription factors (TFs) bind to specific activation site[17], this association can induce long-range chromatin loop when the promoter and enhancer are brought together transiently [18]. Interestingly, there are also proteins such as remodelers that constantly bind to open or close chromatin. If genes are likely to be expressed in open DNA regions, it remains unclear how a region can stay open after TFs leave their binding regions.

According to present analysis, when a chromatin loop is formed, through possible loop extrusion or due to the local proximity of chromatin induced by TFs and remodelers, this situation facilitates the binding and rebinding of cross-linkers, such as diffusing CTCF molecules, which can attach and detach DNA dynamically. Such a dynamical binding and unbinding can lead to a preferential stabilization of a region where promoter-activator have initially generated a transient loop. This situation is equivalent to positive feedback process, where a loop is stabilized in a region enrich in cross-linkers. Since CTCF molecules could be trapped in this nanoregion, these molecules have less chance to diffuse away. These molecules enter and leave this nanoregion and their presence maintains loops.

This elementary mechanism guarantees the presence of cross-linkers at the base of a loop, since each cross-linker that could escape is replaced by others entering this local region. This dynamical mechanism keeps the initial TF-remodeler chromatin changes and thus stabilizes a loop, initially induced during a specific gene activation. Possibly such mechanism could be responsible for a certain fraction of loops. Actually, cohesin and CTCF are not always involved in loop formation [19] and loops could be formed by sporadic encounters of distant DNA sequences [9, 20].

The present mechanism is not related to the mechanism of loop formation, such as the loop extrusion model [21], where loops are generated by an active process, increasing the loop length until cohesin stops at the boundary made by two CTCF sites with the right orientation. Here, on the contrary, we consider that a loop is already formed and we studied the conditions for its stability. Finally, when the Ω-loop domains are sufficiently separated, the present analysis can be generalized to the entire nucleus. In a nucleus containing around 10^3^ TADs [4], there are 10^4^ loops[22], 10^5^ cohesin and 10^5^ CTCF molecules (concentration ~ 144.3 nM) [23]. We hypothesize that if there are in average ~ 10 CTCF molecules per TAD, as single cell Hi-C has revealed, then we are left with few loops per TADs [24].

In this final section, we propose to estimate the mean life of chromatin loops based on the present model and the binding *λ* and unbinding *μ* rate constants for CTCF and cohesin molecules. We will use the experimental data provided in [25]. Using the resident unbinding time 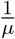 which is given by ~ 1 − 2*min* and ~ 22*min* for CTCF and cohesin respectively. In addition, the binding time (reciprocal of the rate *λ*) is given by ~ 1*min* and ~ 33*min* for CTCF and cohesin respectively. As mentioned above, using that there are in average *N* = 10 cross-linkers (CTCF and cohesin molecules) per loop, we shall now provide some estimates for the loop stability time, which is the mean passage time of the last resident time of the cross-linker to the base of a loop formula summarized by formulas 4 and 8 according to the first and the second scenarios respectively. For the case CTCF molecules, using the rates 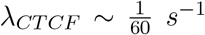 and the range 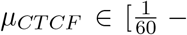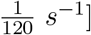, we get for stability time 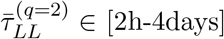,(*q* = 2 because we need at least 2 CTCF to create one loop) for the first scenario. For cohesin with *N* = 10 molecules per loop and the rates 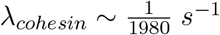 and 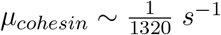, we obtain a stability time of 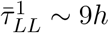(*q* = 1 because one cohesin can forma loop).

We shall now explore the possibility to have a heterogeneity distribution of molecules per loop: when there are only 2 CTCF molecules, the stability time reduced to 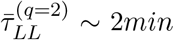), instead of the 2 hours. However when increasing *N* = 10 to *N* = 12, the stability time increases to 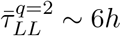. To conclude, the present model suggest that chromatin sites with only two CTCF molecules support loops that unstable of the order of minutes, in contrast with average sites with 10 and more molecules, for which loops are stable for hours. Similarly, the presence of only two cohesin rings is associated with a stability time of 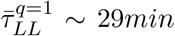 (instead of 9 hours with 10 molecules). By increasing *N* to 12, this leads to 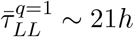. To conclude, one or two cohesin are associated with a stability of the order of tens of minutes, while 10 molecules leads to a stability of few hours.

In summary, in this manuscript, we presented possible scenarios to study the chromatin organization by stabilizing loops, depending on the local numbers of CTCF and cohesins. We hypothesized that these number allows to keep the loop for minutes to days (memory hypothesis). In our scenario, the presence of at least one molecule allows the loop binding domain to exit and thus further molecules can bing this domain, leading to a positive feedback of binding cross-linkers (fig. 7A-C). More specifically, in this memory hypothesis, initially a local transient chromatin loop is induced by promoter-enhancer contacts. The local proximity of chromatin induced by transcription factors and remodelers facilitates the binding of looping factors such as the diffusing cohesin molecules, which can attach and detach DNA dynamically. This dynamical binding and unbinding of cross-linker molecules leads to a preferential stabilization of loops, which result in keeping the memory of the initial TF-remodeler chromatin change. However, if such a dynamical balance between binding and unbinding cross-linker is disrupted, the local loop organization of chromatin is lost. This memory mechanism hypothesis should be further explored and tested experimentally. Finally, the present model suggests that there is a spectrum of loop stability, ranging from few minutes to the entire cell cycle depending on the number of cross-linkers per loop site.

**FIG. 7.**
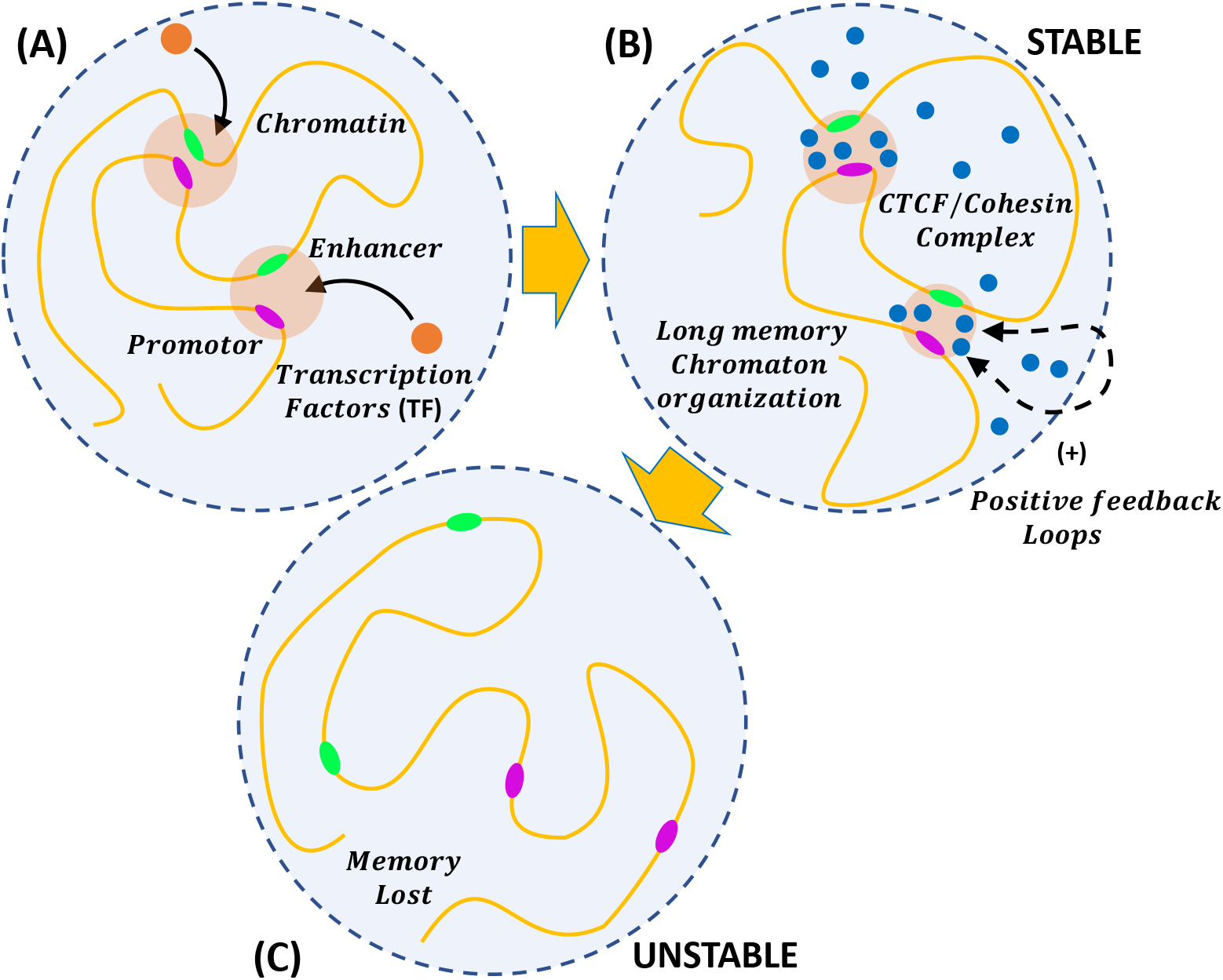
Chromatin memory hypothesis induced by positive feedback binding. **A** Local transient chromatin loop is induced by promoter-enhancer contacts thus making various loops. The local proximity of chromatin induced by transcription factors (TF) and remodelers facilitates the binding of looping factors such as the diffusing cohesin molecules, which can attach and detach DNA dynamically. **B** Dynamical binding and unbinding of cohesin molecules lead to a preferential stabilization of loops through a positive feedback process which keeps the memory of the initial TF-remodeler chromatin change. **C** If the dynamical balance between binding and unbinding cohesin molecules is not disrupted, the local loop organization of chromatin is lost.

## APPENDIX

We present in the appendix the computation related to the three cases.

## EXPLICIT COMPUTATION FOR MODEL 1

Using eq. 3, we rewrite the system of differential equations 2 using the initial conditions *p*_*k*_ = *δ*_*kq*_ and the limit of *t* → ∞: *p*_0_ (∞) = 1 and *p*_*k*_ (∞) = 0. We get:

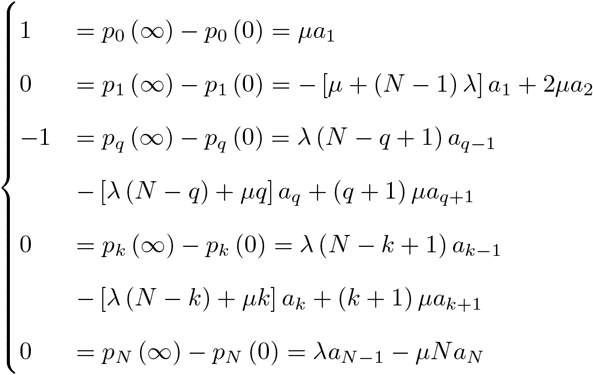

We compute here the sum 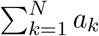 with respect to the parameters *λ*, *μ* and *N*. The equations involving *a*_1_ and *a*_2_ resolve as

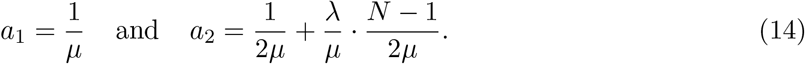

In general, we obtain the induction relation:

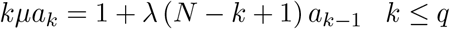

while

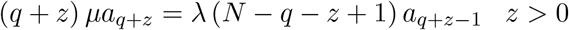

We express the last equation in term of *a*_*q*_:

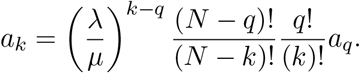

where

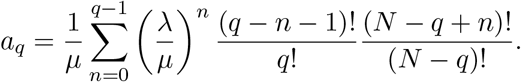

These relations are summarized as

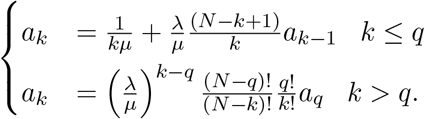

We split the sum for the expression of 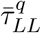 according to the initial number *q* of cross-linkers and we get

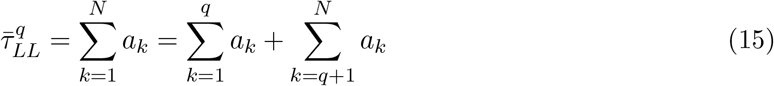

where

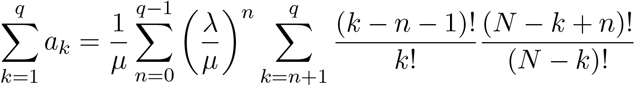

and

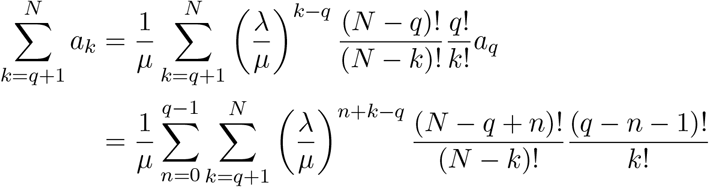

Finally, we obtain the general expression

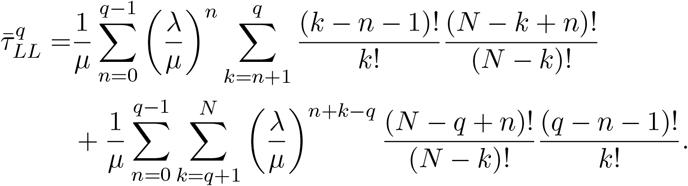

## EXPLICIT COMPUTATION FOR MODEL 2

Following the method of induction described in the previous section of the appendix (expressions as eq. 14), we obtain the induction relation for *a*_*k*_ for the second scenario when the there are initially *q* bound cross-linkers:

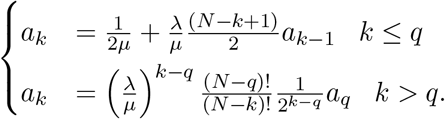

Using relation 15, we compute the time 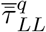 using the three relations

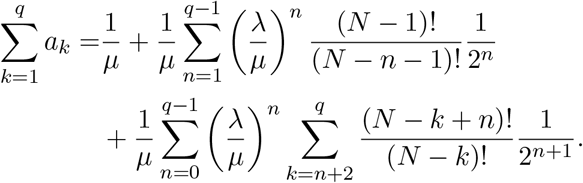

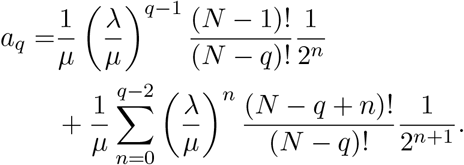

and the sum for *k* > *q*:

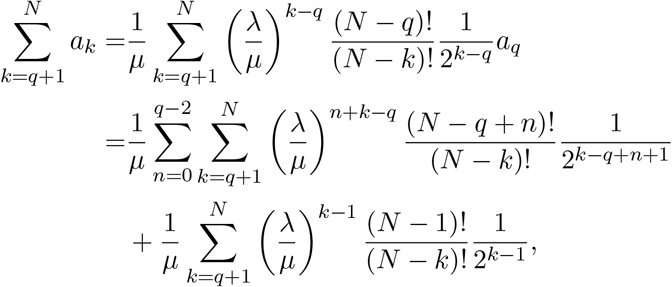

leading to the general expression:

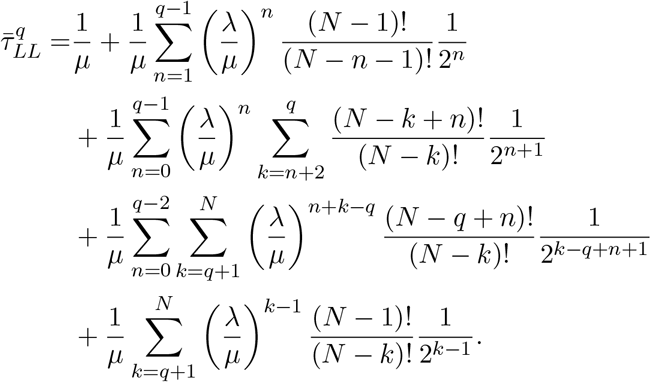

## EXPLICIT COMPUTATION FOR MODEL 3

In the third scenario, the steady-state coefficients *a*_1_ and *a*_2_ are again similar to the ones of equations 14 while the induction relation for *a*_*k*_ becomes:

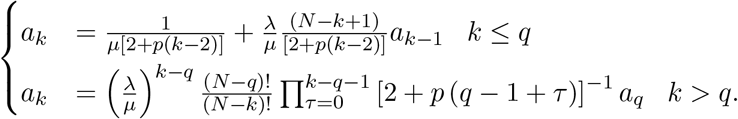

To evaluate the time 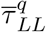, we use the sums

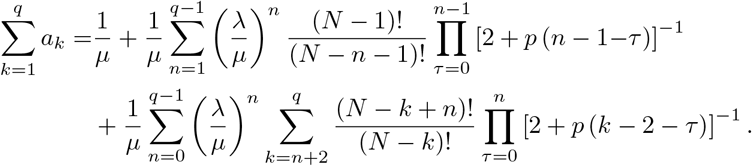

Since

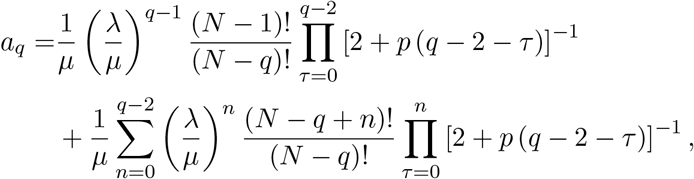

we can obtain the sum for *k* > *q*:

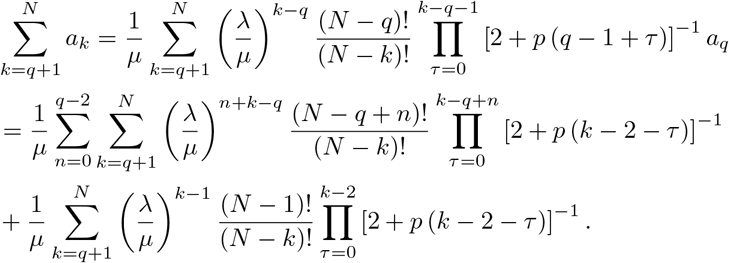

Finally, we obtain the general expression is:

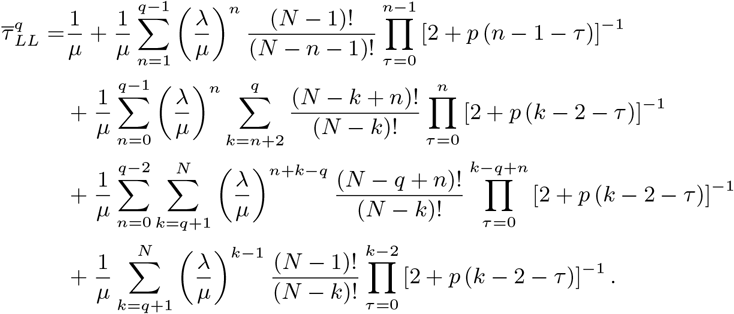

## ACKNOWLEDGEMENTS

This work was supported by Plan Cancer and Fondation pour la Recherche Médicale (FRM)-[SPF201909009284]. This project has received funding from the European Research Council (ERC) under the European Unions Horizon 2020 research and innovation programme (grant agreement No 882673).

## AVAILABILITY OF DATA

The analysis (Wolfram Mathematica sheets) that supports the figures of this work are available from the corresponding author upon reasonable request.

